# RiVIERA-beta: Joint Bayesian inference of risk variants and tissue-specific epigenomic enrichments across multiple complex human diseases

**DOI:** 10.1101/059329

**Authors:** Yue Li, Manolis Kellis

## Abstract

Genome wide association studies (GWAS) provide a powerful approach for uncovering disease-associated variants in human, but fine-mapping the causal variants remains a challenge. This is partly remedied by prioritization of disease-associated variants that overlap GWAS-enriched epigenomic annotations. Here, we introduce a new Bayesian model RiVIERA-beta (<u>Ri</u>sk <u>V</u>ariant <u>I</u>nference using <u>E</u>pigenomic <u>R</u>eference <u>A</u>nnotations) for inference of driver variants by modelling summary statistics p-values in Beta density function across multiple traits using hundreds of epigenomic annotations. In simulation, RiVIERA-beta promising power in detecting causal variants and causal annotations, the multi-trait joint inference further improved the detection power. We applied RiVIERA-beta to model the existing GWAS summary statistics of 9 autoimmune diseases and Schizophrenia by jointly harnessing the potential causal enrichments among 848 tissue-specific epigenomics annotations from ENCODE/Roadmap consortium covering 127 cell/tissue types and 8 major epigenomic marks. RiVIERA-beta identified meaningful tissue-specific enrichments for enhancer regions defined by H3K4me1 and H3K27ac for Blood T-Cell specifically in the 9 autoimmune diseases and Brain-specific enhancer activities exclusively in Schizophrenia. Moreover, the variants from the 95% credible sets exhibited high conservation and enrichments for GTEx whole-blood eQTLs located within transcription-factor-binding-sites and DNA-hypersensitive-sites. Furthermore, joint modeling the nine immune traits by simultaneously inferring and exploiting the underlying epigenomic correlation between traits further improved the functional enrichments compared to single-trait models.

## 1 Introduction

Genome wide association studies (GWAS) can help gain numerous insights on the genetic basis of complex diseases, and ultimately contribute to personalized risk prediction and precision medicine [1–4]. However, fine-mapping the exact causal variants is challenging due to linkage disequilibrium (LD) and the lack of ability to interpret the function of noncoding variants, which contribute to about 90% of the current GWAS catalog (40.7% intergenic and 48.6% intronic; [5]). On the other hand, several lines of evidence have been proposed to help interpret non-coding genetic signals, in order to gain insights into potential regulatory functions. In particular, epigenomic annotations can pinpoint locations of biochemical activity indicative of cis-regulatory functions [6,7]. Indeed, comparison with genome-wide annotations of putative regulatory elements has shown enrichment of GWAS variants in enhancer-associated histone modifications, regions of open chromatin, and conserved noncoding elements [3,6,8–12], indicating they may play gene-regulatory roles. These enrichments have been used to predict relevant cell types and non-coding annotations for specific traits [6,9,13]. Furthermore, many complex traits potentially share causal mechanisms such as autoimmune diseases [14,15] and psychiatric disorders [16,17]. Thus, methods that jointly model the intrinsic comorbidity implicated in the GWAS summary statistics of the related traits may confer higher statistical power of causal variants detection. Recently, several methods were developed to utilize the wealth of genome-wide annotations primarily provided by ENCODE consortium to predict causal variants and novel risk variants that are weakly associated in complex traits. Pickrell (2014) developed a statistical approach called fgwas that models association statistics of a given trait and used regularized logistic function to simultaneously learn the relevant annotations. To account for LD, fgwas assumes at most one causal variants per locus via a softmax function. Kichaev et al. (2014) recently developed a multivariate Gaussian framework called PAINTOR, which allows for more than one causal SNP but at most three to be located within a single locus by considering all of the combinatorial settings [18]. Chung et al. (2014) used a maximum likelihood framework called GPA to infer driver variants shared among multiple traits by modeling the corresponding GWAS p-values as Beta distributions with an option of using one or more sets of annotations to improve the power detecting causal variants [19]. Although useful, these methods are often designed to simultaneously operate on a small number of independent annotations due to some computational constraints. Moreover, most methods only operate on one trait at a time whereas exploiting the correlation between traits at the epigenomic annotation level may prove useful for shared causal mechanisms that go beyond the level of individual variants.

In this article, we describe a novel Bayesian framework called RiVIERA-beta (<u>Ri</u>sk <u>V</u>ariant Inference using <u>E</u>pigenomic <u>R</u>eference <u>A</u>nnotations to model <u>Beta</u> likelihood of GWAS summary statistics p-values). The main novelty of RiVIERA-beta is the ability to perform efficient Bayesian inference of the intrinsic causal signals across multiple traits while simultaneously inferring and exploiting enrichment signals and their correlation between traits over hundreds of tissue-specific epigenomic annotations. We achieve this efficiently via stochastic sampling of loci and powerful Hamiltonian Monte Carlo sampling of model parameters [20]. We first use simulation to demonstrate the utility of RiVIERA-beta in prioritizing driver variants and detecting functional epigenomic annotations. We then apply RiVIERA-beta to some of the most well-powered GWAS datasets, consisting of 9 immunological disorders from ImmunoBase [15] and Schizophrenia 2014 data from Psychiatric Genomic Consortium [21]. To infer tissue-specific epigenomic enrichments, we utilize the largest compendium of epigenomic annotations to date from ENCODE/Roadmap Consortia, consisting of 848 annotations including 8 major epigenomic marks across 127 distinct cell types [7]. This allows us to revisit the GWAS of these 10 common complex disorders by inferring their underlying regulatory variants implicated at the tissue-specific epigenomic contexts.

## 2 MATERIALS AND METHODS

### GWAS summary statistics

The GWAS summary statistics for the nine immune diseases were obtained from ImmunoBase (March 17, 2015) [15]. The nine diseases are: Autoimmune Thyroid Disease (ATD), Celiac Disease (CEL), Juvenile Idiopathic Arthritis (JIA), Multiple Sclerosis (MS), Narcolepsy (NAR), Primary Biliary Cirrhosis (PBC), Psoriasis (PSO), Rheumatoid Arthritis (RA), Type 1 Diabetes (T1D). We imputed the p-values of un-genotyped SNPs using FAPI and 1000 Genome European data (Phase 1 version 3) [22]. We then obtained the p-values of SNPs that fall within the pre-defined risk loci available from ImmunoBase for each of the 9 immune traits. For all analyses, we filtered out risk loci or variants in the MHC regions or sex chromosomes X and Y. The Schizophrenia 2014 (SCZ2) summary data containing 642846 observed and imputed SNPs were obtained from Psychiatric Genomic Consortium (PGC) [21]. Among these, 54132 SNPs fall within the 105 SCZ-associated loci of the autosomes (chr 1-22) defined by PGC (we filtered out the 3 loci on chromosome X). **Table 1** summarizes the total number of SNPs and risk loci for each individual GWAS that were subject to the proposed fine-mapping analyses.

**Table 1:** GWAS data summary

| Abbrev | Trait | Total | Loci | gwSNPs | cSNP_st | cSNPs_mt |
| --- | --- | --- | --- | --- | --- | --- |
| ATD | Autoimmune Thyroid Disease | 4206 | 8 | 630 | 38 | 49 |
| CEL | Celiac Disease | 29784 | 39 | 2592 | 344 | 211 |
| JIA | Juvenile Idiopathic Arthritis | 13427 | 22 | 3 | 440 | 223 |
| MS | Multiple Sclerosis | 61360 | 104 | 2096 | 884 | 339 |
| NAR | Narcolepsy | 1316 | 3 | 62 | 22 | 16 |
| PBC | Primary Biliary Cirrhosis | 14573 | 19 | 2498 | 172 | 111 |
| PSO | Psoriasis | 24832 | 34 | 457 | 305 | 171 |
| RA | Rheumatoid Arthritis | 38207 | 78 | 470 | 1978 | 719 |
| SCZ2 | Schizophrenia | 54132 | 105 | 5217 | 2481 | NA |
| T1D | Type 1 Diabetes | 41945 | 57 | 2832 | 826 | 327 |

We investigated 10 GWAS traits as listed above. Abbrev: abbreviation of the trait names; Total: total number of SNPs in the risk loci with imputed and observed summary statistics; Loci: total number of risk loci for each trait; gwSNPs: SNPs that pass GWAS cutoff $p < 5e-8$ ; cSNP\_st: total number of SNPs that are included into the 95% credible set based on single-trait risk inference using RiVIERA-beta; cSNP\_mt: SNPs in 95% credible set constructed based on multi-trait joint risk inference using RiVIERA-beta across the 9 immune traits (without SCZ2).

### Roadmap epigenome data

Roadmap epigenome data were obtained from Roadmap epigenomic web portal (March, 2015). Peaks were defined if their p-values were below 0.01 (i.e., following the definition of “Narrow Peaks” [7]). In total, there are 848 epigenome tracks, including 8 epigenomic marks namely H3K4me1, H3K4me3, H3K36me3, H3K27me3, H3K9me3, H3K27ac, H3K9ac and DNase I in in 127 cell or tissue types, which were grouped into 19 categories [7]. To associate each SNP with the annotations, we overlapped their genomic coordinates with each bigWig epigenome track making use of the R packages *rtracklayer* and *GenomicAlignments*. SNPs that fall within a peak of an annotations will have value 1 otherwise 0 for that annotation. The resulting matrix is a *V_d_* × *K* input matrix containing the epigenomic values across *K* = 848 marks for each of the *V_d_* SNPs in disease *d*.

### Running existing fine-mapping software on simulated data

#### fgwas

The software `fgwas` [23] (version 0.3.4) were downloaded from GitHub. We prepared the input for fgwas (1) the Z scores calculated as the t-statistics of the linear coefficients of the genotype of each variant fitted separately by least square regression on the simulated continuous phenotypes (**Materials and methods**) and (2) 100 discretized epigenomic annotations at p < 0.01. To enable fine-mapping, we issued -`fine` flag and specify the region numbers for each SNP in the input file as required by the software. As part of the outputs from fgwas, we obtained ‘PPA’ and ‘estimate’ for the causal variants and influences of each epigenomic annotations, respectively.

#### GPA

GPA (0.9-3) [19] was downloaded from GitHub and run with default settings. Same as above, we set the annotations to one at p-value < 0.01 and 0 otherwise. To test for trait-relevant annotations, we followed the package vignette. Briefly, we fit two GPA models with and without the annotation and compared the two models by `aTest` function from GPA, which performs likelihood-ratio (LR) test via *χ*^2^ approximation, and obtained the enrichment scores as the -log10 p-value.

#### PAINTOR

PAINTOR (version 2.1) was downloaded from GitHub [18]. As suggested in the documentation, we prepared a list of input files for every locus including summary statistics as t-statistics, LD matrices, and binary epigenomic annotations. We ran the software with default setting with assumption of at most two causal variants per locus. We then extracted the ‘Posterior_Prob’ and ‘Enrichment.Values’ as the model predictions for causal variants and causal annotations, respectively.

### Details of RiVIERA-beta Bayesian model

#### Inference of empirical prior *π*_*vd*_

We first define the empirical prior function of a variant *v* being associated with disease *d* as a logistic function:

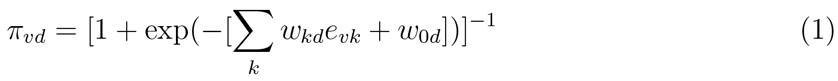

where *w_kd_* ∈ **w**_*d*_ denotes the linear coefficient or the influence of the *k^th^* epigenomic mark affecting disease *d* and *w*_0*d*_ is the linear bias.

We assume that epigenomic causal effect *w_kd_* follows a multivariate Gaussian distribution with zero mean and unknown covariance:

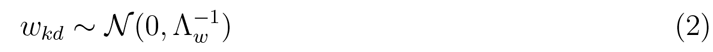

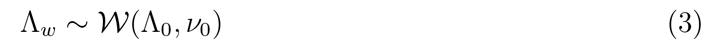

where Λ_*w*_ is a *D* × *D* inverse covariance matrix 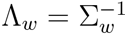 to model the pairwise epigenomic correlation among *D* diseases. It follows a Wishart distribution with identity matrix as prior (i.e., by default, we assume *aprior* no correlation between the target traits) and *v*_0_ = 0 (i.e., by default, we did observe any samples *aprior* that are indicative of the correlation between any two diseases being modeled). The hyperparameters can be easily modified to incorporate prior belief on the correlation between any two diseases of interests.

Additionally, the bias *w*_0*d*_ follows a Gaussian distribution with unknown variance and mean determined based on our prior belief of the causal fraction *π*_0_:

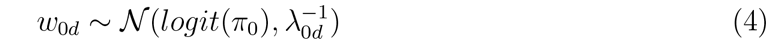

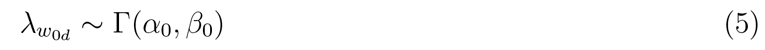

where 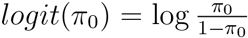. By default, we set *π*_0_ to 0.01, implying that 1% of the SNPs in the risk loci are expected to be causal when no functional enrichment. We set *α* = 0.01 and *β* = 0.0001 to enable a broad hyperprior for *w*_0*d*_.

Notably, *w_kd_* can be interpreted as enrichment coefficient for annotation *k* in disease *d*, where a positive *w_kd_* will increase the causal prior *π*_*vd*_ when *e_vk_* = 1. During the training, however, w_kd_ may become negative, which makes the interpretation difficult. Thus, we constrain *w_kd_* to be non-negative values, which involves imposing infinitely high potential energy for negative *w_kd_*. More details are described in **Supplementary Text 1**.

#### Inference of variant causality *c_vd_* given prior *π*_*vd*_ and model parameters *μ_d_, ϕ_d_*

Because the target association variable *a_vd_* for variant *v* in disease *d* represents p-values, which are continuous and restricted to the interval (0, 1), we assume that it follows a Beta distribution with unknown mean *μ_d_* and unknown precision *ϕ_d_*:

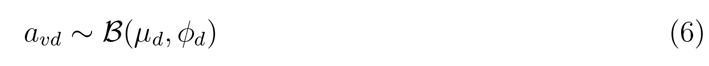

Note that we re-parameterize Beta density function from the traditional “rate” *p* and “shape” *q* parameters, and instead use mean *μ* = *p*/(*p* + *q*) and precision *ϕ* = *p* + *q*, as per [24, 25]. Specifically, the density function of association variable *a_vd_* is defined as follows:

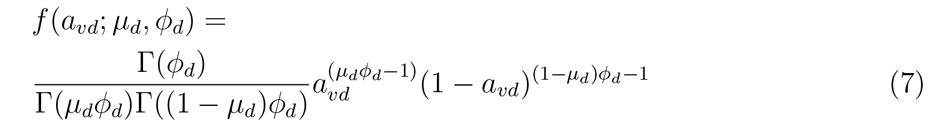

Further, we let the mean *μ_d_* and precision *ϕ_d_* follow Beta and uniform prior, respectively:

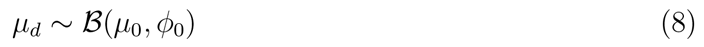

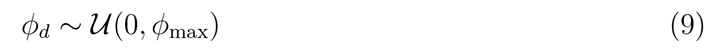

where the hyperparameters (*μ*_0_, *ϕ*_0_) reflect *apriori* belief on the p-value signal of a causal variant. By default, we set *μ*_0_ = 0.1 and *ϕ*_0_ = 2. If *ϕ*_max_ = ∞, *ϕ* follows an improper prior. Because it is unlikely to have a very large *ϕ*, by default, we set *ϕ*_max_ to 1000. Notably, as long as *ϕ*_max_ is large, the inference results remain the same with different *ϕ*_max_ values.

With the prior *p*(*c_vd_*|**w**_*d*_, **e**_*v*_) ≡ *π*_*vd*_ and likelihood *p*(*a_vd_*|*μ_d_*,*ϕ_d_*) ≡ *f*(*a_vd_*; *μ_d_,ϕ_d_*) established, the posterior probability of association (PPA) [26] of variant *v* being causal for disease *d* then follows:

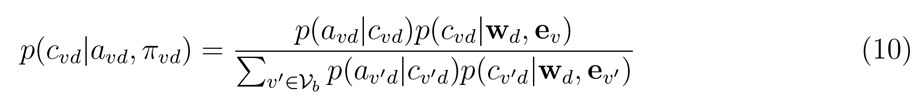

where 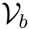 represent all variants within locus *b*. The 95% credible set 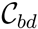 for each locus *b* is the minimal number of SNPs 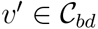 in the locus such that 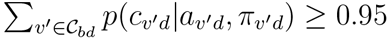.

#### Joint posterior distribution

The complete likelihood density function treating *c_vd_* as missing values is defined as:

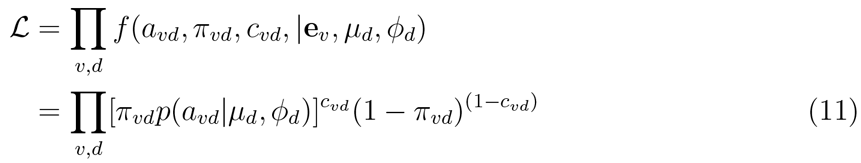

The logarithmic joint posterior density function is then:

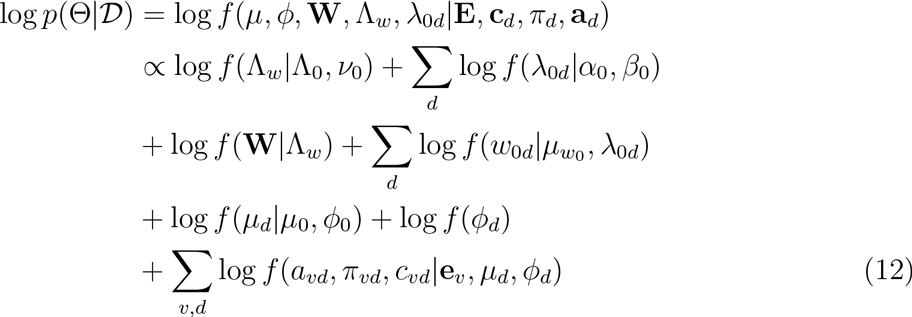

In principle, causality is inferred by integrating out all nuisance parameters:

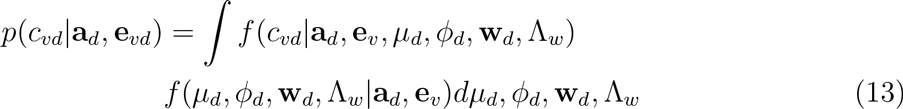

which is not tractable. We employ Markov Chain Monte Carlo (MCMC) to sample from the joint posterior.

#### Markov Chain Monte Carlo

We use Gibbs sampling [27] to sample the precision matrix Λ_*w*_ of epigenomic effects from the posterior distribution. Specifically, Gibbs sampling requires a closed form posterior distribution. Due to the conjugacy of the Wishart prior of epigenomic precision Λ_*w*_ to the multivariate normal distribution of epigenomic effect **W**, the posterior of the epigenomic precision matrix Λ_*w*_ also follows Wishart distribution [28]:

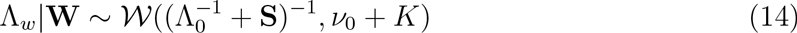

where **S** is the sample variance of **W**, i.e., *S* = **W^*T*^W**.

Similarly, we sample *λ*_0*d*_ from Gamma posterior distribution:

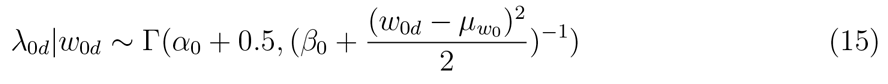

To sample epigenomic effects **w**_*d*_, prior bias *w*_0*d*_, causal mean *μ_d_*, causal precision *ϕ_d_* for disease *d* = 1, &, *D*, we employ a more powerful gradient-based sampling scheme namely Hamiltonian Monte Carlo (also known as hybrid Monte Carlo) (HMC) [20,29], exploiting the fact that the joint posterior of our model is differentiable with respect to the model parameters *μ_d_,ϕ_d_,w_kd_,w_0d_* (**Supplementary Text S1**). Finally, after discarding *t*% models accepted before the burn-in period (default: t=20%), we obtain the Bayesian estimates of PPA by averaging the corresponding values computed over the *T′* individual models accepted throughout the *T* MCMC runs.

#### Bayesian fold-enrichment tests for epigenomic annotations

Due to co-linearity among the epigenomic annotations, directly using *w_kd_* to assess the epigenomic enrichment for annotation *k* may be misleading. We propose an heuristic approach to assess the log fold-enrichment of the full prior model over the alternative prior with the effect of annotation *k* for disease *d* removed (i.e., **w**_*d\k*_,*w_kd_* = 0):

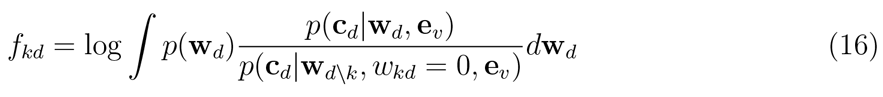

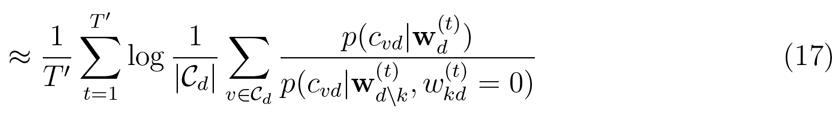

where 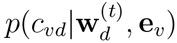 is the logistic prior based on Eq 1, 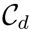 is the union of all the 95% credible sets across loci for disease *d*: 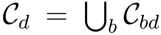 Notably, under the optional constraint that *w_kd_* ≥ 0, *f_kd_* is always positive, which directly translates to fold-enrichment of annotation *k* conditioned on all the other annotations *k′* ≠ *k*. The 95% Bayesian credible interval for *f_kd_* are obtained from the *T′* MCMC runs. The significance of each annotation *k* is determined based on the ranking of its lower bound *f_kd_* (i.e., the 2.5% quantile of *f_kd_*).

Alternatively, we can estimate the fold-enrichment for each annotation simply based on the ratio of estimated fraction of causal variants in an annotation *e_vk_* over the fraction of all of the variants in that annotation 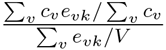, where *c_v_* is the PPA for SNP *v*. This is more efficient and accurate when the underlying causal variants were randomly sampled from the annotations as done in the simulation.

#### Stochastic gradient updates per locus

Directly updating model parameters based on the gradients of all GWAS loci at each MCMC iteration is inefficient and results in poor HMC acceptance rate. Instead, at each MCMC update, we randomly sample one locus and update the model parameters (which are shared across loci) based on that locus. We find this approach quite efficient in capturing meaningful causal properties such as causal signals and relevant epigenomes that are shared across all risk loci. Together, we outline the overall algorithm of the proposed Bayesian model in Algorithm S2 (**Supplementary Text S1**).

### GWAS simulation

To assess the power of the proposed fine-mapping model in identifying causal variants and compare it with existing methods, we implemented a simulation pipeline adapted from [18]. Briefly, the simulation can be divided into three stages (1) simulate genotypes based on the haplotypes from 1000 Genome European data (phase 1 version 3) using HapGen2 [30] (**Supplementary Fig. S1**); (2) simulate epigenomic enrichments and subsequently sample causal variants accordingly using 100 Roadmap annotations selected from each of the 19 categories of primary tissue/cell types (**Supplementary Fig. S1**); (3) simulate liability phenotype plus the random noise to obtain the desired heritability (fixed at 0.25) and subsequently the GWAS summary statistics in terms p-values and z-scores via ordinary least square regression. Details are described in **Supplementary Text**.

### Gene ontology enrichment analysis

We obtained the latest gene annotations from Ensembl database (version 80) programmatically via biomaRt package [31], which resulted in 10,801 gene ontology (GO) terms in biological processes (BP). To assign SNPs to genes, we performed lift-over to map the SNPs from hg19 to hg38 using rtracklayer [32] and assigned each SNP to a gene if it is located within 35 kb up and 10 kb downstream of that gene. The resulting Ensemble gene identifiers were matched with those genes in each GO-BP category. We then performed hypergeometric tests on each GO-BP term for all of potential *in-cis* target genes of the SNPs in each trait and adjusted for multiple testings using Benjamini-Hocherg family-wise Type I error correction method [33]. For the 9 immune traits, the enrichment signals are strong so we set the cutoff at FDR < 0.005; for Schizophrenia, we set FDR < 0.2.

### RiVIERA-beta software

RiVIERA-beta is available as an open-source R package with documented functions and walk-through examples described in the vignette. Most functions were implemented in C++ by integrating *Rcpp* and *RcppArmadillo* libraries [34]. These libraries enabled us to apply RiVIERA-beta to large matrices very efficiently with complied code and having much lesser memory overhead than a naïe R implementation. RiVIERA-beta is available at Github (https://github.mit.edu/liyue/rivieraBeta).

## 3 RESULTS

### RiVIERA model overview

The fundamental hypothesis of our model is that non-coding disease associations are driven by disruption of regulatory elements of common activity patterns (e.g., motifs of sequence-specific regulators), thus leading to gene expression changes and ultimately phenotypic changes at the cellular or organism level between case and control individuals. Our RiVIERA-beta Bayesian model aims to infer the probability that a given variant *v* is a driver for disease *d* by modeling the corresponding GWAS association statistic for that variant using a vector of genome-wide epigenomic annotations (**e**_*v*_). Given a set of *B* risk loci, the inputs to RiVIERA-beta are GWAS summary statistics in terms of p-values and a set of discrete or continuous epigenomic annotations (**Fig**. 1a). In this study, we used binary signals to ease interpretation of the functional enrichments. We train RiVIERA-beta by repeatedly sampling one locus at each iteration to efficiently learn the intrinsic (i.e., locus-independent) causal signals. **Fig**. 1b depicts RiVIERA-beta as probabilistic graphical model [35]. The observed variable of our model is the GWAS association values (in terms of p-values) *a_vd_*xs for each variant *v* in each disease *d*. We assume that *a_vd_* follows a Beta distribution with unknown mean and dispersion parameters. The effect of each annotation on each trait is learned as global annotation-by-disease weight matrix **w**, which follows a *D*-dimensional multivariate normal distribution with zero mean and *D* × *D* disease-disease covariance Λ_w_. The prior probability *π*_*vd*_ that a variant *v* is causal in disease *d* is essentially a linear combination of the weighted genomic annotations **e**_*v*_, which reflects the disease-associated active histone marks and DNA accessibility in the 127 cell types (**Materials and methods**). The outputs of the model (**Fig**. 1c) are (a) posterior probability of association (PPA) *c_vd_* that variant *v* is causal in disease *d*; (b) the Bayesian fold-enrichment estimates *f_kd_* based on the ratio between the full prior model with all annotations over the null prior model with all annotations except for annotation *k*.

**Figure 1:**
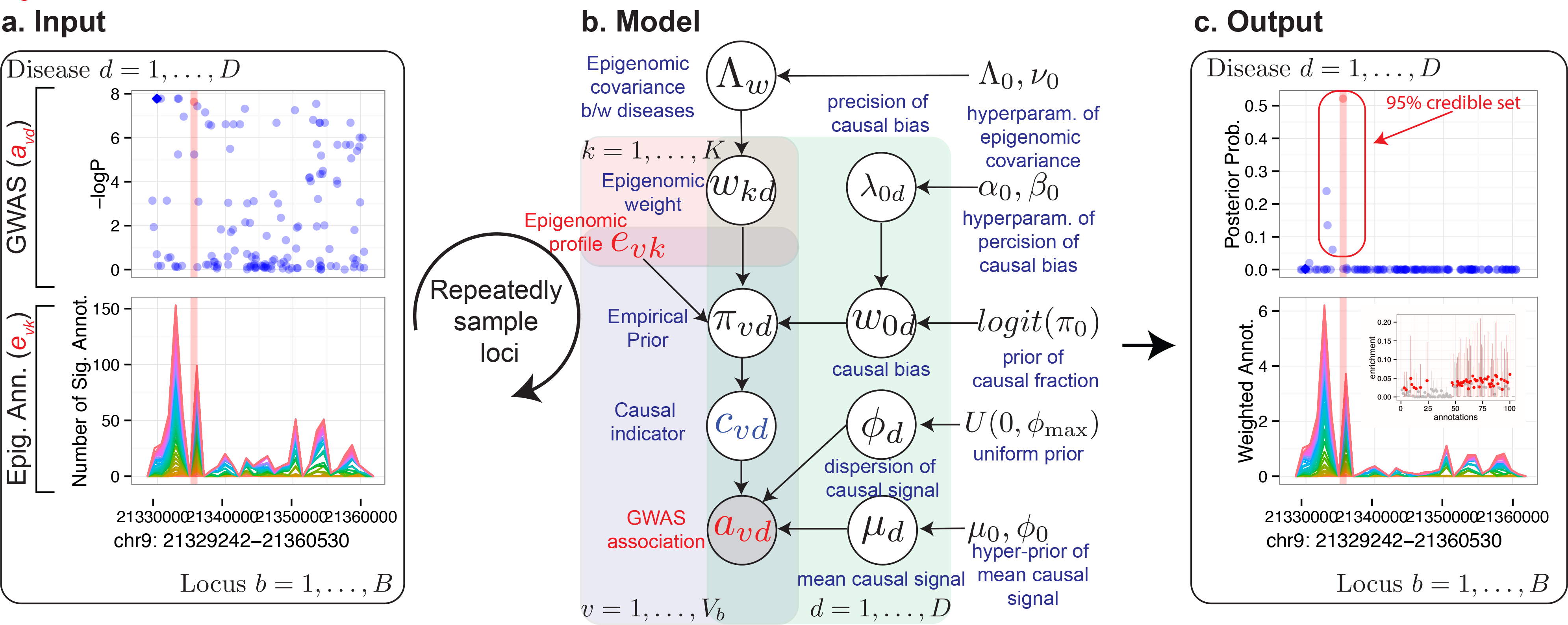
RiVIERA-beta model overview. **(a)** Inputs to RiVIERA-beta are GWAS summary statistics and epigenomic annotations for *B* risk loci. At a given iteration, the model samples one locus and tries to learn the intrinsic causal signals implicated in the corresponding GWAS summary data and epigenomic profiles. Highlighted variant is the causal variant based on the simulated data. **(b)** The probabilistic graphical model representation of RiVIERA-beta [35]. Variables for which distribution is defined are in circle. Epigenomic profiles are treated as observed values with no circle. The variable in shaded circle are observed (i.e., GWAS association *a_vd_* and variables in unshaded circle are unobserved. The variables in red are observed and variables in blue are the variables of interest (i.e., causal indicator). The two colored plates represent *K* annotations (red) and *V* variants (blue). We model the GWAS association *a_vd_* of variant *v* in terms of p-value sampled from Beta distribution with unknown precision *ϕ_d_* and mean *μ_d_*, which respectively follow an uninformative prior and a Beta distribution with hyperparameters *μ*_0_, *ϕ*_0_. The latent variable *c_vd_* indicates whether variant *v* is causal in disease *d*. On top of it, we dedicate an empirical prior as a linear combination of the epigenomic profile *e_vk_* weighted by the epigenomic influence *w_kd_*, which follows multivariate normal with zero mean and a *D* × *D* inverse covariance or precision matrix 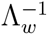, where *D* is the number of traits that are being modeled. The linear bias *w*_0*d*_ expresses the prior belief of the causal fraction *π*_0_ (default: 0.01). **(c)** Outputs from the model are posterior probabilities of association (PPA) for each variant in each locus, the 95% credible set containing the minimal number of SNPs whose PPA sum to 0.95 or greater, and the Bayesian estimates of the fold-enrichment of each annotation.

### Method comparison using GWAS simulation

The goal of the simulation is to evaluate the model’s power to predict (1) causal variants in each locus; (2) the relevant annotations that determine which variants are causal. To this end, we simulated GWAS summary statistics based on 1000 Genome European data (Phase 1 release 3) (**Supplementary Fig. S1**) and 100 representative epigenomic annotations (**Supplementary Fig. S1**) (**(Materials and methods**). We performed a series of power analyses over 500 simulation runs.

First, we examined how well the posterior probabilities were calibrated by taking the credible SNPs that contribute to 95% posterior mass inferred by each method (**Supplementary Fig. S2**). As expected, when our model assumption of single-causal variant per locus holds, we observe that our model is well calibrated (**Fig**. S2), where the 95% credible SNPs indeed correspond to approximately 95% of the causal variants. When there are more than one causal variants per locus, the 95% credible SNPs include on average 50% the true causal SNPs (**Supplementary Fig. S2**)

Because the number of variants within the credible set differs depending on the concentration of the posterior probabilities inferred by each method, we sought to control that bias by evaluating the proportion of identified causal variants as a function of the absolute number of selected variants. When the assumption of one-causal-variant-per-locus holds, we observed comparable or better performance of RiVIERA-beta compared to existing methods (**Fig**. 2). As expected, when the assumption is violated, our current model is second to PAINTOR, which is able to infer multiple causal variants per locus (**Supplementary Fig. S3**). We also examined the correlation between the functional enrichments estimated by each method and the underlying epigenomic enrichments that were used to simulate the causal variants. The performance of the four methods are comparable with the proposed model achieving a slightly better correlation (**Supplementary Fig. S4**).

**Figure 2:**
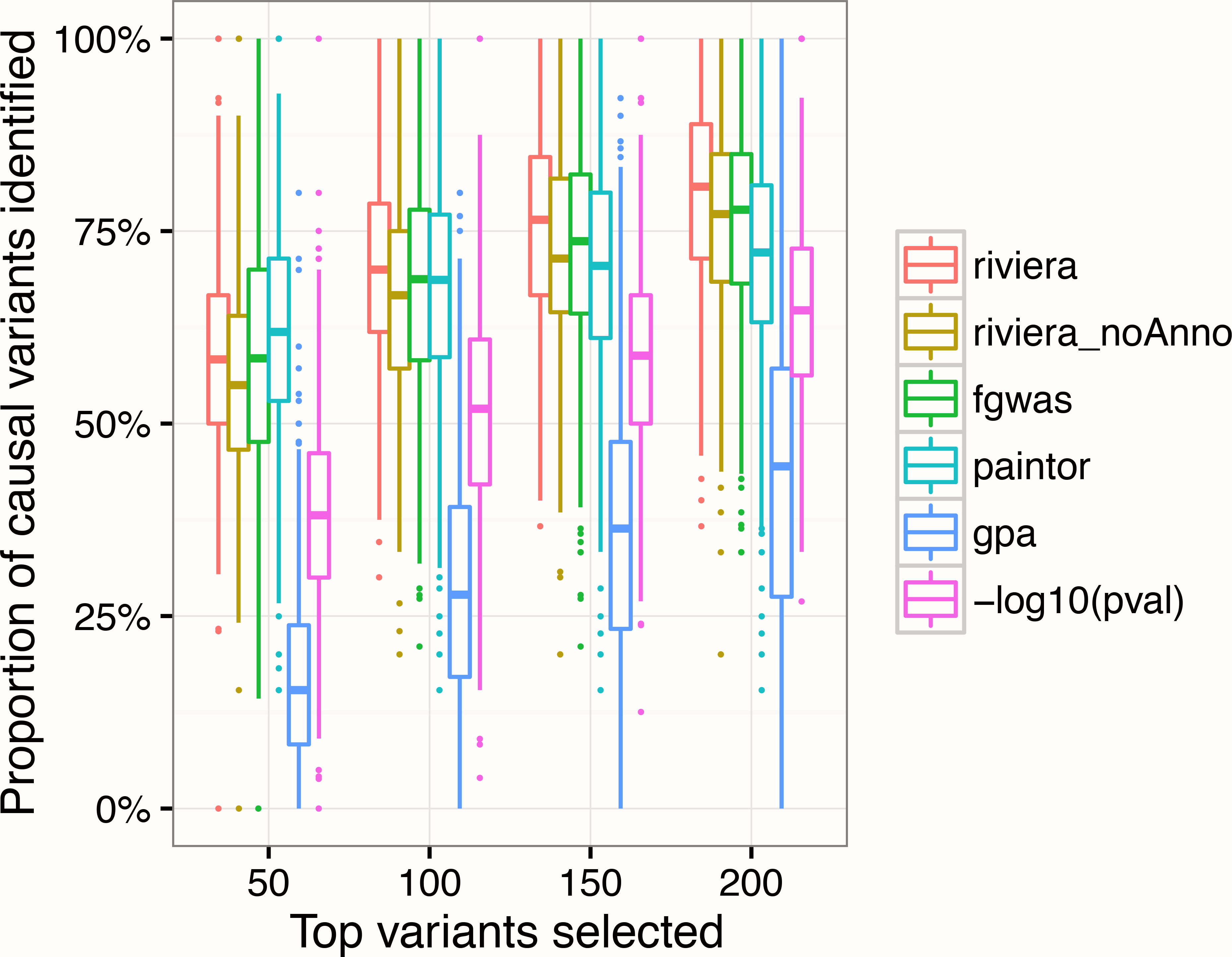
Model performance on simulated datasets. Proportion of causal variants were identified by each method as a function of increasing number of top variants selected.

### Applications to immune and psychiatric disorders

To demonstrate RiVIERA-beta in a real-world application, we used it to investigate 10 complex diseases including 9 immune diseases with summary statistics obtained from ImmunoBase [15] and Schizophrenia from Psychiatric Genomic Consortium (PGC) [21] (Table 1) We used 848 epigenomic annotations from ENCODE/Roadmap consortium (**Materials and methods**) to build a functional prior for each trait to aid fine-mapping and conduct cell-type specific epigenomic enrichment analyses [7]. We first applied RiVIERA-beta to the 10 traits separately to examine individual causal signals and then demonstrated RiVIERA-beta’s capability to operate on the 9 immune traits and the improved detection power compared to the single-trait model.

### RiVIERA-beta detected meaningful tissue-specific enhancers in test GWAS traits

We first sought to confirm the validity of the model through its ability to identify meaningful cell-types or tissues for each trait. To this end, we selected the top 5% (i.e., the top 43) of the 848 annotations for each disease based on the corresponding Bayesian estimates of the lower bounds of the 95% credible interval (**Supplementary Table S1**; **Materials and methods**). We then performed hypergeometric tests on enrichments of each of the 19 categories grouped by Roadmap consortium based on the cell types and tissues [7]. Indeed, we observed a significant enrichment for Blood & T-cell for all 9 immune disorders but not for Schizophrenia, which exhibits exclusive epigenomic enrichments in the Brain category (Hypergeometric adjusted p-values < 0.05) (**Fig**. 3a). Additionally, we also observed modest enrichments for B-cell and Thymus tissue in the 9 immune traits. We then examined the enrichment status for the 8 epigenomic marks. Indeed, enhancer marks namely H3me4me1 and/or H3K27ac are most significantly enriched among all 8 marks (q-values < 0.05). In addition, H3K4me3 associated with promoter is also enriched in most immune traits. Interestingly, we also observed a modest enrichment of H3K9me3 in Schizophrenia but not in the immune traits. We further ascertained the enrichment results by re-running RiVIERA-beta on the permuted data matrix and observed diminishment of the meaningful enrichment observed above (**Supplementary Fig. S5**).

**Figure 3:**
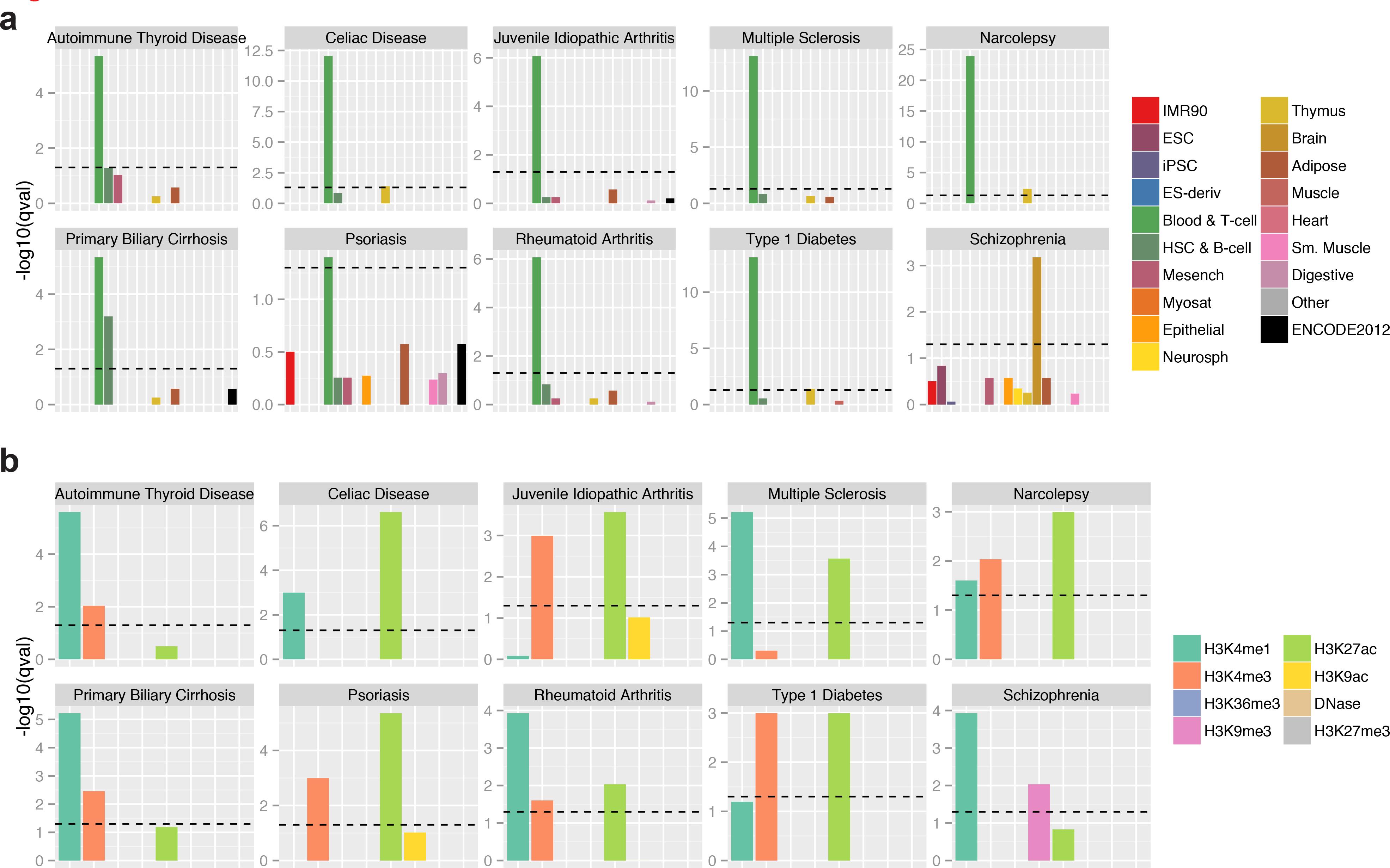
Predicted tissue-specific epigenomic enrichments in the 10 GWAS traits. (**a**). Hypergeometric enrichment for each of the 19 primary tissue categories using the top 5% or 43 annotations out of the 848 annotations in total for each trait based on the lower bound of the 95% credible interval of the Bayesian fold-enrichment estimates by our RiVIERA-beta; model; (**b**) enrichments for the 8 epigenomic marks among the top 43 annotations for each trait. Y-axis is the logarithmic q-values, which are the corrected p-values from the hypergeometric tests for multiple testing across traits and tissue groups or marks by Benjamini-Hochberg method [33]. On both plots, horizontal dashed bars indicate standard statistical threshold of FDR < 0.05.

### SNPs in the credible set exhibit promising regulatory potentials

The variants in the credible set are more enriched for functional elements. Inspired by the promising tissue-specific enhancer enrichment results obtained above, we refined our RiVIERA-beta model by re-training it on the top 5% (or 43) annotations on each trait using the same GWAS data. For each locus in each trait, we then constructed 95% credible set (**Supplementary Table S2**; **Materials and methods**). On average, we were able to construct a rather small credible set ranging from 4 to 25 SNPs per locus for the 10 traits (**Table** 1). As a comparison, we extracted the same number of SNPs with the most significant GWAS p-values from each locus. For ease of reference, we named our SNPs in the credible set as “credible SNP” and the GWAS counterpart as “GWAS SNP”. Compared to GWAS SNPs, the credible SNPs exhibit substantially higher averaged placental conservation scores (phastCons46way obtained from UCSC database) across most traits (**Fig**. 4 CONS).

**Figure 4:**
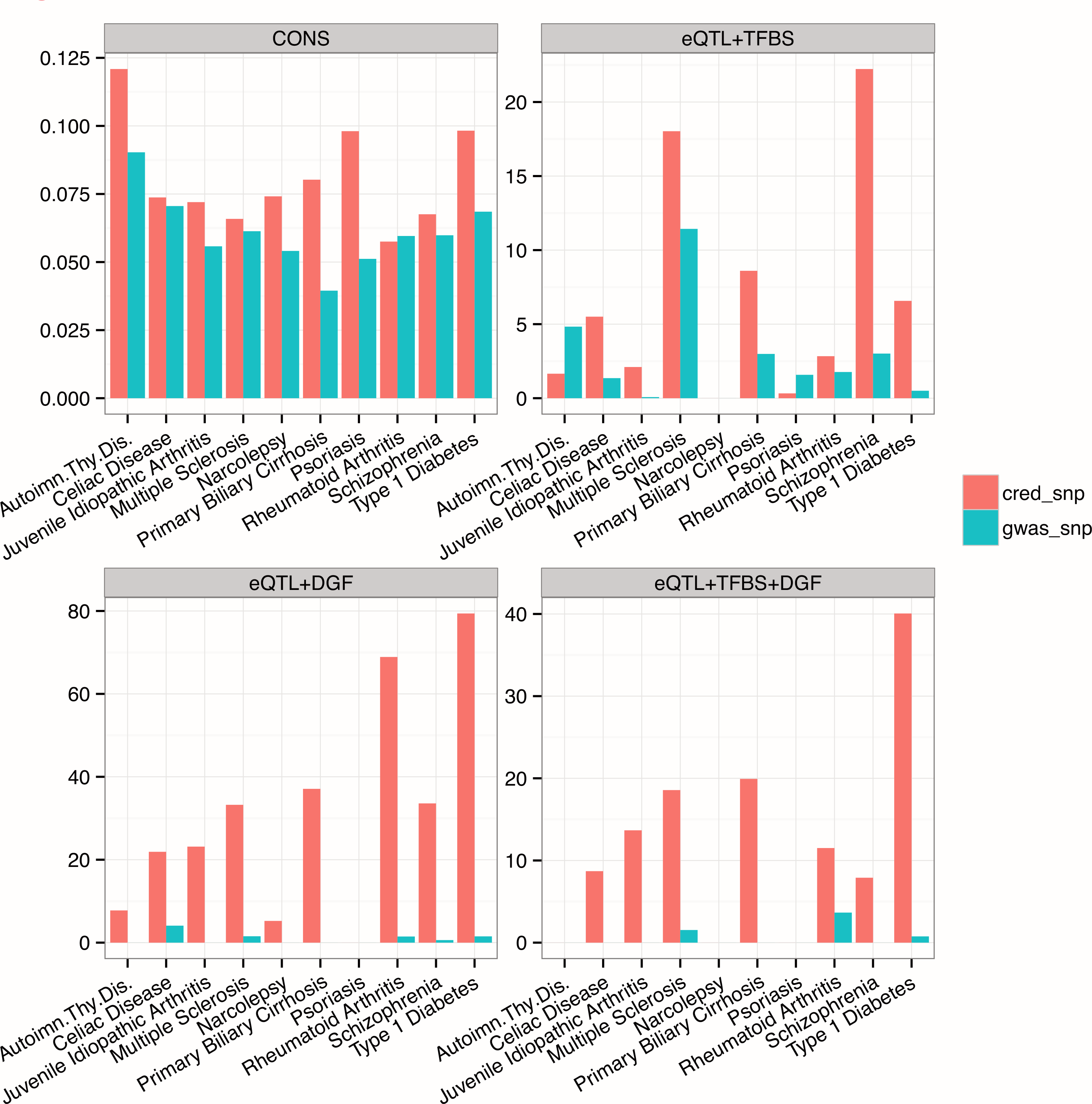
Functional enrichments of credible SNPs. The top left panel displays the averaged phastCons46way conservation scores for variants in the 95% credible set (cred_snp) and the same number of SNPs chosen based on GWAS p-values (gwas_snp). The three other panels illustrate hypergeometric enrichments in terms of the-log10 q-values corrected for multiple testing over the 10 traits of the selected variants for GTEx whole blood eQTL located within transcription factor binding sites based on sequence motif (TFBS) (eQTL+TFBS) and genomic digital footprint (DGF) (eQLT+DGF), and eQTL in both TFBS and DGF (eQTL+TFBS+DGF).

Moreover, the credible SNPs were significantly enriched for expression quantitative trait loci (eQTL) that are in the regulatory regions. Specifically, we obtained in total 806,847 GTEx whole-blood eQTL-SNPs (version 6) [36] and retained 122,549 and 23,973 eQTL-SNPs that overlap with transcription factor binding sites derived from 1,772 TF recognition motifs [37] and digital genomic footprints (DGF) at 6-bp resolution derived from DNaseI data in CD cells using method described in [38], respectively as well as 6,743 eQTL-SNPs that overlapped with both the TFBS and DGF regions. We then performed hypergeometric tests to assess the significance of overlap between the credible/GWAS SNPs and the regulatory-eQTL SNPs. Indeed, our credible SNPs exhibit much higher enrichments for those eQTL-SNPs, suggesting their regulatory potentials elucidated based on the enhancer activities by our proposed RiVIERA-beta model (**Fig**. 4; **Supplementary Table S3**).

### Gene-centric analysis revealed enrichment for meaningful biological processes

Genes adjacent to the SNPs in credible sets are significantly enriched for disease-specific biological processes. In particular, we observed significant enrichments of many immune-related processes for the *in-cis* genes for which the SNPs in the credible set are within 35 kb upstream or 10 kb downstream (**Fig**. 5; **Supplementary Table S4**; **Materials and methods**). For instance, regulation of T cell homeostatic proliferation, regulation of interferon-gamma-mediated signaling pathway, and regulation of type I interferon-mediated signal pathways are among the most significantly enriched GO terms in 5 or 6 out of the 9 immune traits. In contrast, the enrichments for Schizophrenia are dominated by GO terms involving synaptic processes and neuronal differentiation/development. The enrichment results are mostly consistent between the credible genes and the genes derived from the same number of SNPs chosen based on the GWAS p-values (GWAS-genes).

**Figure 5:**
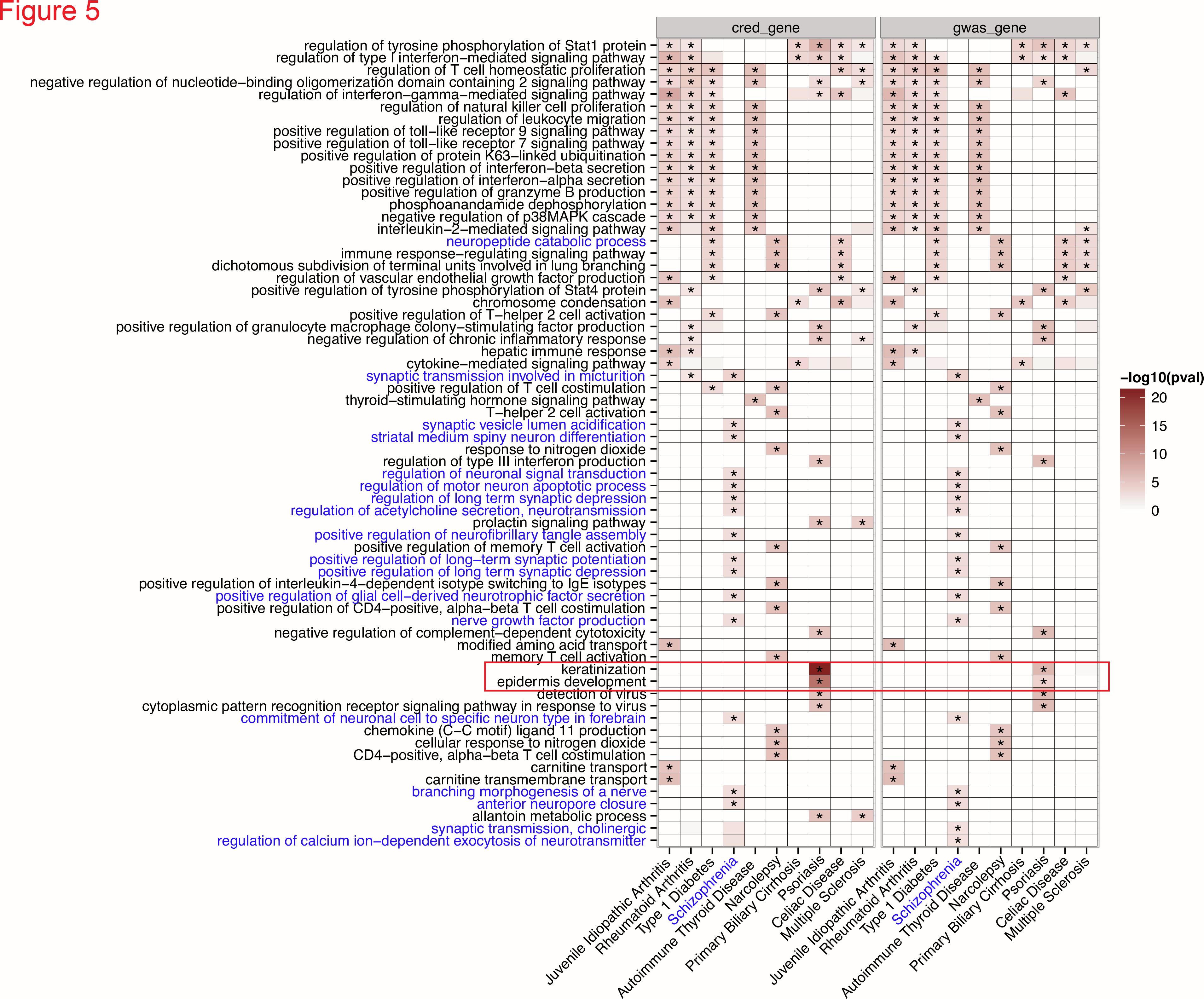
Gene ontology enrichments across the 10 traits. Rows are the GO biological processes and columns are the 10 traits. Color intensities in each cell reflect the significance level in terms of-log10 p-value. Asterisks indicate q-values above significant cutoff after correcting for multiple testings (FDR < 0.2). GO names that match the pattern ‘synap|neuro|nerve’ are colored blue to highlight their exclusive association with ‘Schizophrenia’ (also in blue). Notably, GO terms ‘keratinization’ and ‘epidermis development’ (highlighted in the red box) are exclusively enriched for Psoriasis. Diseases were ordered based on hierarchical clustering based on the Pearson correlation of their GO enrichment scores.

Intriguingly, we observed a highly significant enrichment for keratinization (GO:0031424) and epidermis (e.g., skin) development (GO:0008544) exclusively for Psoriasis. In particular, 17 genes among the 241 credible genes belong to keratinization and epidermis development, which contain in total 49 and 121 genes, respectively (q < 9 × 10^−18^, q < 2 × 10^−10^). Indeed, Psoriasis is mainly characterized as a chronic skin disease with epidermal hyperproliferation [39,40]. In contrast, there are only 6 out of 157 GWAS-genes are defined in each of two GO categories (q < 0.001).

To further ascertain the RiVIERA-beta fine-mapping results, we created a visualization scheme for each of the 469 risk loci across 10 traits examined (**Supplementary Fig. S6**). Fig. 6 displays two example loci for Type 1 diabetes (chr17: 37383069-38239012) and Schizophrenia (chr7: 104598324-105062839). The upper panel displays the RiVIERA-beta model prior, the genetic signals from GWAS-log p-values, and RiVIERA-beta PPA. Red colored and diamond shape points are GTEx whole-blood eQTL SNP and top SNPs included into 60% credible set (we used 60% to not clutter the plot with the remaining SNPs in the 95% credible set that exhibit low PPA). Intuitively, SNPs with high PPA exhibit both prominent genetic and epigenetic signals. Thus, to infer causal variants, RiVIERA-beta efficiently took into account not only the GWAS signals derived from the genetic data but also the prior signals mainly driven by the weighted epigenomic profiles. The middle panel illustrates the cumulative density for each epigenomic profiles weighted by the tissue-specific enrichment estimates.

**Figure 6:**
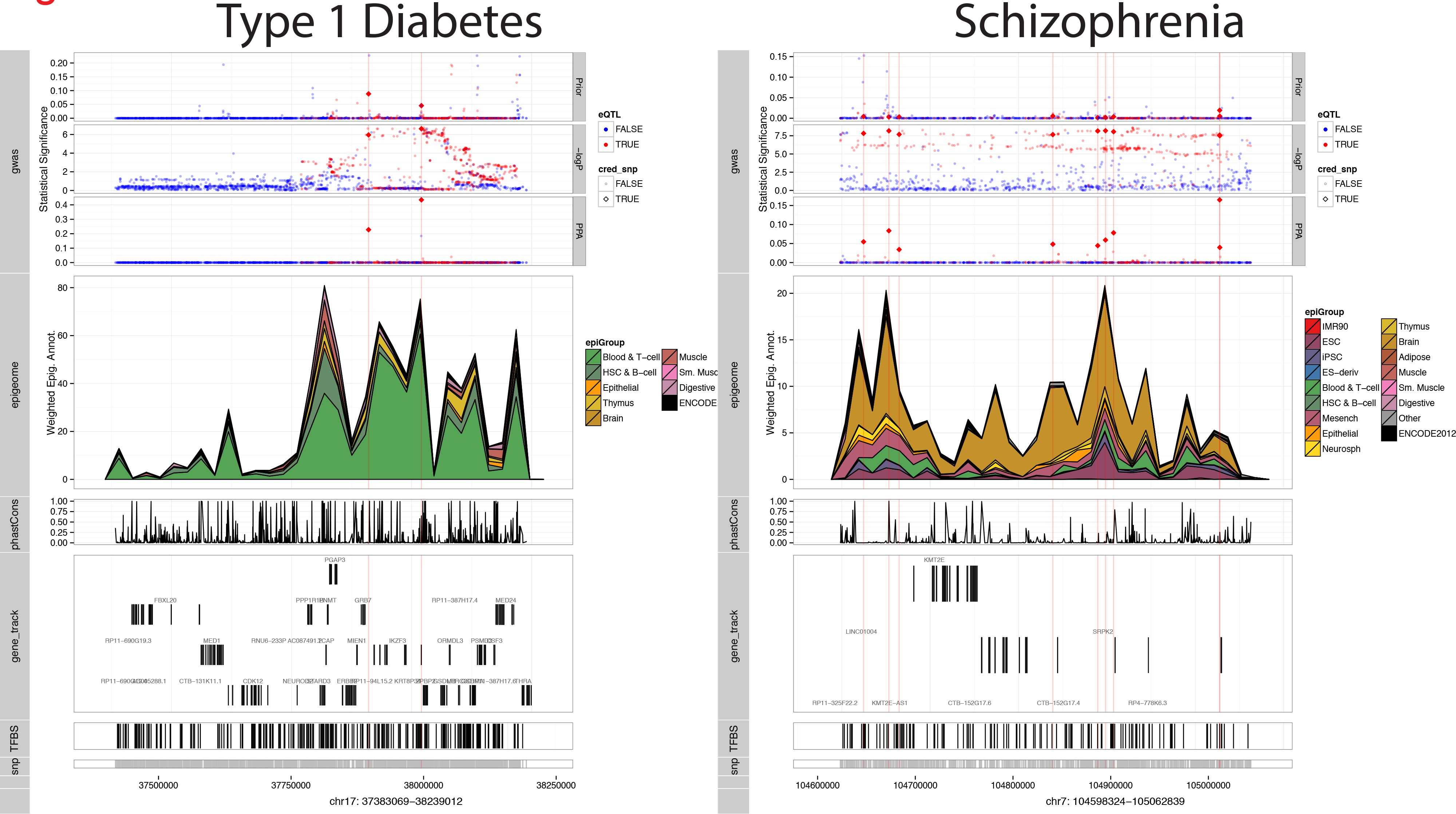
Visualization of fine-mapping results. Top track: the upper panel display the RiVIERA-beta prior, genetic signals of GWAS-log10 p-values, and RiVIERA-beta PPA; the middle track illustrates the cumulative density of weighted epigenomic profiles colored based on the epigenomic group; the bottom tracks shows the conservation, gene annotations (Gencode 19), transcription factor binding sites (TFBS), and SNP positions. The red colored and bigger diamond plots indicate whole-blood GTEx eQTL SNPs and SNPs included into the 60% credible set, respectively. For illustration purpose, only one risk locus for Type 1 diabetes and one for Schizophrenia are shown above. The full display of 469 risk loci were in **Supplementary Fig. S6**.

Consistent with the overall enrichment results (**Fig**. 3) we observed prominent enrichments for the enhancer regions predominantly in blood T cells for all of the 9 immune traits and brain tissue for Schizophrenia. The bottom tracks display the external functional information (i.e., not in the training data) including conservation score, genes, transcription factor binding sites based on motif matches that may further aid variant selection for downstream experimental validation (please refer to **Supplementary Table S2** for detailed information). We also visualized the signals within the of Psoriasis-associated risk region ch1:152536784-152785170, which harbors genes involved in keratinization and epidermis development as mentioned above. Interestingly, as an exception of most other immune-susceptible loci, the underlying epigenomic profiles exhibit prominent signals not only in blood T cell but also in epithelia enhancer regions (**Supplementary Fig. S6**). However, the associated SNPs exhibit rather weak genetic signal perhaps due to lower allele frequencies.

### Multi-trait causal inference improved functional enrichments in most immune traits

Exploiting epigenomic correlation between highly related immune diseases improved functional enrichments in several traits. We performed multi-trait causal inference over all of the 9 autoimmune traits by jointly applying our RiVIERA-beta to 364 risk loci concatenated together from the 9 immune traits using 174 epigenomic annotations which was a union of unique annotations from the top 43 annotations for each individual trait. We focused only on the 9 immune GWAS (i.e., leaving out Schizophrenia) because they commonly utilized the same genotyping platform namely ImmunoChip. The multi-trait GWAS summary statistics triggered RiVIERA-beta to infer the disease covariance matrix and sample disease-specific epigenomic weights from the joint posterior with modified zero-mean multivariate normal prior that takes into account the sampled disease covariance (**Materials and methods**). As a results, RiVIERA-beta sampled correlated epigenomic weights between traits more frequently compared to the single-trait model.

We constructed the 95% credible sets for each trait using the disease-specific PPA derived from the joint model and assessed the functional enrichments as above (**Supplementary Table S6**). Notably, the cross-trait model exploited 174 annotations as apposed to 43 annotations used by the single-trait model. To examine whether the improved enrichments were due to the increased number of annotations, we re-ran a single-trait model for each of the 9 traits separately each using the 174 annotations. Compared to the 95% credible set constructed based on the single-trait causal inference using the top 43 annotations, we observed smaller 95% credible sets for 8 out of the 9 immune traits (**Table** 1), suggesting that the mulit-trait joint inference further improved the model confidence in some of the highly related traits.

More importantly, we observed a much more improved enrichments for the GTEx whole-blood eQTL SNPs located within open chromatin regions or digital genomic footprints in most of the immune traits (**Fig**. 7; **Supplementary Table S5**). Thus, the joint inference further improved the regulatory potential through following the Hamiltonian trajectory that is more consistent with the epigenomic correlation pattern between the related immune traits. We also repeated the GO enrichment analysis on the 95% credible set and found that the enriched GO terms were mostly immune-specific biological processes and consistent with the above single-trait analyses (**Supplementary Fig. S7**; **Supplementary Table S7**).

**Figure 7:**
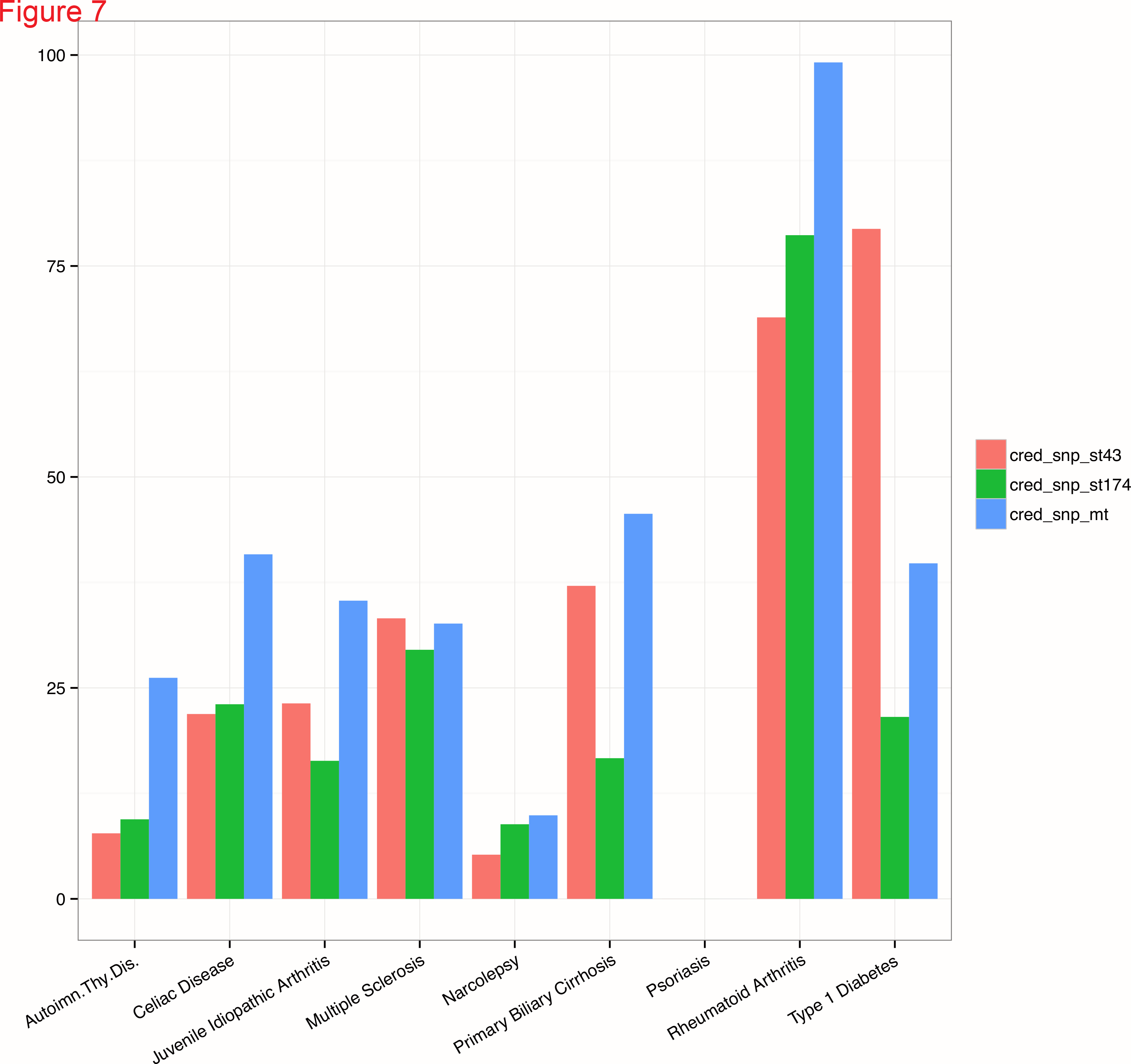
Enrichments for eQLT using credible SNPs constructed from multi-trait joint inference. Credible SNPs for each trait were constructed based on PPA inferred by the joint RiVIERA-beta model over the 9 immune traits using 174 annotations, which are the union of the top 43 annotations detected from each trait individually. We then assessed the hypergeometric enrichments of the 95% credible sets for the GTEx whole-blood eQTL that are within DNA hypersensitive sites as defined by the genomic digital footprint data [38]. We compared these enrichment scores derived from the multi-trait model (cred_snp_mt) to the enrichments derived from the single-trait models either running on 43 annotations (cred_snp_st43) or on the 174 annotations (cred_snp_st174). The latter was included txso control for the improvements due to the increased number of annotations (from 43 to 174).

## 4 DISCUSSION

Understanding the genetic basis of complex traits hinges upon powerful integrative methods to map genotypes to phenotypes [41]. Fine-mapping causal variants has been a highly active and fruitful research in the past several years [9,18,42–44]. However, most existing methods typically operate solely on genetic data by estimating each SNP of being causal conditioned on the lead index SNPs in the same LD block, which are typically approximated by the 1000 Genome data [9,15,45,46]. With the recent availability of large-scale functional genomic data provide by ENCODE/Roadmap consortia, there is an urgent need to incorporate these valuable information in a principled way as a form of Bayesian prior. In this article, we describe a novel Bayesian fine-mapping method RiVIERA-beta to re-prioritize GWAS summary statistics based on their epigenomic contexts. The main contribution of RiVIERA-beta is the ability to systematically construct from a targeted set of susceptible loci a Bayesian credible set of SNPs, which exhibit plausible tissue-specific regulatory properties implicated in the large epigenomic data compendium either in a single trait or across multiple traits.

One benefit of using the raw epigenomic annotations rather than using the inferred signals such as ChromHMM [7] or GenoSkyline [47] states derived from the annotations is that it eases the interpretation of the actual relevant epigenomic marks in the relevant tissue types and facilitates downstream experimental efforts to assay the specific marks in the disease-specific cell types. However, the correlation of the epigenomic marks will make difficult estimating the underlying functional enrichments. Moreover, we choose to model the summary statistics rather than genotypes because it is not always possible to obtain individual-level phenotype-genotype data particularly for large-scale meta-analysis. Thus, effective methods based on summary statistics may benefit wider research communities than methods that solely operate on individual-level genotype data [18,19,23]. Moreover, our model *requires only p-values* because it uses Beta distribution to model the likelihood. In contrast, fgwas requires both the z-scores and the standard error from the linear regression used in the GWAS to estimate the Wakefield approximate Bayes factors. While some recent GWAS summary statistics include those information, there are many do not have z-scores and/or standard error of the linear model but only p-values (e.g., the ImmunoChip data we used in our studies for the 9 immune traits). When the standard error is not available in a given GWAS summary statistics, fgwas needs to be estimate it from the minor allele frequency of a reference panel such as 1000 Genome, which may not be accurate depending the study cohorts. Additionally, modeling p-values via Beta density function only has more relaxing model assumption than modeling z-scores via normal density although both methods are highly effective in practice.

Overall, SNPs included into the credible set exhibit both significant GWAS signal and high prior. In some cases, however, SNPs that were added to the credible set in each locus do not exhibit significant GWAS p-values (**Supplementary Table S2,S6**). This generally occurs when the genetic signals in those loci are weak relative to the SNPs in other loci for the same trait, and the model functional prior eventually dominates the SNP prioritization. Thus, we recommend considering these variants cautiously when designing downstream experiments.

One important assumption of our model is that there is one causal variant per locus, which is reflected by the normalization of variants within each locus so that they sum to 1 in order to obtain PPA and construct 95% credible sets [23]. When this assumption holds, the posterior probabilities are well calibrated (**Supplementary Fig. S2**). However, as demonstrated in our simulation, when this assumption is violated, the PPA is not well calibrated (**Supplementary Fig. S2,S3**). Other existing method such as PAINTOR [18] and CAVIAR [48] employ multivariate normal distribution to model all of the variants within a locus using LD reference panel estimated from 1000 Genome data as the covariance matrix, which allows inferring more than one causal variants per locus. While CAVIER used only summary statistics, PAINTOR is able to employ functional annotations to aid fine-mapping. Both methods require computing the likelihood density across a combinatorial set of causal configurations and therefore still needs to assume at most an arbitrarily small number of causal variants, typically below 10 causal SNPs per locus.

As future works, we will explore potential ways to enable efficient inference of more than one causal variants per locus. Furthermore, we will also explore the potential gain of incorporating trans-ethnic data, which was effectively demonstrated by the trans-ethnic version of the PAINTOR model [49]. Moreover, in addition to modeling the epigenomic correlation between traits, variant prioritization may further benefit by jointly inferring the comorbidity at the individual SNP level [19], gene level [50], and/or pathway level [17]. Together, we believe that RiVIERA-beta will serve as a valuable tool complementary to the existing methods in identifying novel risk variants through tissue-specific epigenome-aware fine-mapping as well as aiding the selection of the appropriate cell types and epigenomic marks for more focused investigations of the disruptions of chromatin states by the disease-specific causal variants.

## 5 ACKNOWLEDGEMENTS

We thank Yongjin Park, Gerald Quon, Abhishek Sarkar, and Zhizhuo Zhang for the helpful discussions.

## Supplementary Data

Supplementary Data are available at NAR Online: Supplementary Text S1,S2; Supplementary Tables S1-S5; Supplementary Figures S1-S6.

## Funding

National Institutes of Health (NIH) [R01-HG004037, RC1-HG005334, R01-HG008155]. Funding for open access charge: NIH [R01 HG004037].

### Conict of interest statement

None declared.

